# Differential Effector Response of Amnion- and Adipose-Derived Mesenchymal Stem Cells to Inflammation; Implications for Intradiscal Therapy

**DOI:** 10.1101/523001

**Authors:** Ryan Borem, Allison Madeline, Mackenzie Bowman, Sanjitpal Gill, John Tokish, Jeremy Mercuri

**Affiliations:** Laboratory of Orthopaedic Tissue Regeneration & Orthobiologics, Department of Bioengineering, Clemson University, Clemson, SC 29634, USA; Department of Orthopaedic Surgery, Medical Group of the Carolinas-Pelham, Spartanburg Regional Healthcare System, Greer, SC, 29651, USA; Department of Orthopaedic Surgery, Mayo Clinic, Phoenix, AZ, 85054, USA

**Author notes:** **Corresponding Author:** Jeremy J. Mercuri, 401 Rhodes Engineering Research Center, Clemson University, Clemson, SC 29634, USA.

**Keywords:** Mesenchymal stem cell, intervertebral disc degeneration, amnion, adipose, inflammation, differentiation

## Abstract

Intervertebral disc degeneration (IVDD) is a progressive condition marked by inflammation and tissue destruction. The effector functions of mesenchymal stem cells (MSCs) make them an attractive therapy for patients with IVDD. While several sources of MSCs exist, the optimal choice for use in the inflamed IVD remains a significant question. Adipose (AD)- and amnion (AM)-derived MSCs have several advantages compared to other sources, however, no study has directly compared the impact of IVDD inflammation on their effector functions. Human MSCs were cultured in media with or without supplementation of interleukin-1β and tumor necrosis factor-α at concentrations produced by IVDD cells. MSC proliferation and production of pro- and anti-inflammatory cytokines were quantified following 24- and 48-hours of culture. Additionally, the osteogenic and chondrogenic potential of AD- and AM-MSCs was characterized via histology and biochemical analysis following 28 days of culture. In inflammatory culture, AM-MSCs produced significantly more anti-inflammatory IL-10 (p=0.004) and larger chondrogenic pellets (p=0.04) with greater percent area staining positively for glycosaminoglycan (p<0.001) compared to AD-MSCs. Conversely, AD-MSCs proliferated more resulting in higher cell numbers (p=0.048) and produced higher concentrations of pro-inflammatory cytokines PGE_2_ (p=0.030) and IL-1β (p=0.010) compared to AM-MSCs. Additionally, AD-MSCs produced more mineralized matrix (p<0.001) compared to AM-MSCs. These findings begin to inform researchers and clinicians as to which MSC source may be optimal for different IVD therapies including those that may promote regeneration or fusion. Further study is warranted evaluating these cells in systems which recapitulate the nutrient- and oxygen-deprived environment of the degenerate IVD.

## INTRODUCTION

Intervertebral disc (IVD) degeneration imparts significant socioeconomic burden resulting in annual direct costs estimated to exceed $100B in the U.S.[1] This pathology is mediated in part by increased levels of pro-inflammatory cytokines including interleukin-1 beta (IL-1β) and tumor necrosis factor – alpha (TNF-α).[2] Elevated levels of these cytokines stimulate resident IVD cells to produce extracellular matrix (ECM) degrading proteases that break down tissue.[3] IVD tissues are unable to regenerate as they have limited blood supply and relatively low numbers of active resident cells.[4] Current treatments for IVD degeneration are not curative; they primarily attempt to alleviate pain and inflammation. Ultimately, this pathology becomes debilitating and thus warrant spinal fusion or IVD arthroplasty. Due to the shortcomings associated with current surgical treatments, the orthopaedic community has turned their attention towards the therapeutic potential of mesenchymal stem cells (MSCs).

Populations of MSCs are found throughout adult tissues; however, their numbers are limited. Thus, MSCs are often harvested and expanded *in vitro* prior to administration for therapeutic purposes. The most common tissue sources of MSCs utilized include bone marrow (BM), adipose (AD), and more recently amniotic membrane (AM). It has been established through laboratory studies that MSCs possess several effector functions which make them excellent candidates as biologic therapies for IVD degeneration. MSCs secrete soluble chemicals (i.e., cytokines, chemokines, and growth factors) including prostaglandin E2 (PGE_2_), transforming growth factor-beta (TGF-β), and TNF-α stimulated gene/protein 6 (TSG-6) that can serve an anti-inflammatory function.[5–7] Additionally, soluble signals released by MSCs have been shown to stimulate resident cells to produce new ECM.[8] Moreover, MSCs have been shown to differentiate into several musculoskeletal cell types and generate new IVD ECM *in vitro*.[9] These findings have prompted clinical trials evaluating the safety and efficacy of intradiscal administration of MSCs to attenuate progression of IVD degeneration. These trials have yielded improved pain and functional outcomes in patients with Pfirrmann grade II-V IVD degeneration. [10,11] However, identification of an MSC source which demonstrates optimal therapeutic outcomes within the complex pathological environment of the IVD has yet to be defined. [8,9,11]

Research has been conducted to compare the differentiation capacity of several MSC sources using established *in vitro* protocols.[12–14] Topoluk et al. aimed to identify an optimal MSC source for bone and cartilage regeneration by comparing the osteogenic and chondrogenic differentiation capacity of AM- and AD-MSCs.[12] These two MSC sources were chosen for study as they have higher yields and impart less donor site morbidity at harvest compared to BM-MSCs. The authors demonstrated that AM-MSCs exhibited enhanced gene and ECM markers for bone and cartilage formation compared to AD-MSCs.[12] More recently, the same authors also demonstrated that AM-MSCs are more effective at chondroprotection and skewing pro-inflammatory M1 macrophages towards a pro-regenerative M2 phenotype compared to AD-MSCs.[15] However, considering MSCs are often administered into the inflamed environment of the degenerate IVD, it is imperative to understand the influence of inflammation on the therapeutic effector functions of AM- and AD-MSCs. Such information will help further identify which of these MSC types may be more optimal for therapeutic use in IVD degeneration. Thus, the objective herein was to determine and compare the effect of inflammation on AM- and AD-MSC proliferation, production of cytokines and differentiation.

## MATERIALS & METHODS

### MSC Expansion

Human AD-MSCs were purchased from Invitrogen. Human AM-MSCs were isolated from via informed consent under an IRB approved protocol from an ethics committee according to previously published methods.[12] All MSCs were expanded under standard culture conditions (37°C with 5% CO_2_) until passage 3 (P3). Culture medium consisted of Dulbecco’s Modified Eagle’s Medium (DMEM) containing 10% fetal bovine serum (FBS), and 1% antibiotic/antimitotic (Ab/Am).

### Impact of Inflammation of MSC Proliferation and Cytokine Production Non-Inflammatory and IVD degeneration-mimetic Inflammatory (INF) Culture Conditions

Non-inflammatory (control) conditions included culturing MSCs in DMEM containing 2% FBS and 1% Ab. Inflammatory (INF) media consisted of culture media described above supplemented with human recombinant IL-1β (500pg/ml; PeproTech) and TNF-α (400pg/ml; PeproTech). The concentration of the proinflammatory cytokines used was chosen to mimic levels produced within degenerate (e.g., Pfirrmann Grade IV-V) human IVD tissues. [16] Cells were seeded at 1 × 10^5^ into tissue culture-treated 12-well plates (n=3/condition/cell-type) and cultured for 12- and 48-hours.

### MSC Proliferation

Proliferation of MSCs was evaluated by determining the total cell count per well (n=3 wells/treatment/time-point) using a TC20 Automated Cell Counter (Bio-Rad).

### Cytokine Array and ELISA Analysis of Cell Culture Media for MSC Cytokine Production Studies

Media samples (n=3/condition/cell-type/time-point) were analyzed for several inflammatory cytokines including pro-inflammatory interleukins-1 beta (IL-1β), −1 alpha (IL-1α), −6 (IL-6), −8 (IL-8), and monocyte chemotactic protein-1 (MCP-1) as well as anti-inflammatory interleukins −4 (IL-4) and −10 (IL-10), using a quantitative glass slide array (RayBiotech) according to manufacturer’s instructions. Pro-inflammatory prostaglandin E2 (PGE_2_) was also analyzed from media samples (n=3/condition/cell-type/time-point) via enzyme-linked immunosorbent assay (Abcam) according to the manufacturer’s instructions. All values were calculated as the difference in cytokine concentration (pg/ml) produced by MSCs relative to media only controls.

### Impact of Inflammation on MSC Osteogenesis and Chondrogenesis Non-Inflammatory and Inflammatory Osteogenic Culture Conditions

To induce osteogenic differentiation, 1.0 x10^5^ cells were seeded in 12-well plates (n=3/condition/cell-type) and cultured in monolayer in differentiation media (StemPro Osteogenic Media; ThermoFisher Scientific) for 28 days. For inflammatory conditions, differentiation media was supplemented with IL-1β and TNF-α at concentrations described above.

### Analysis of ECM Mineralization

Osteogenic differentiation was assessed histologically via Alizarin Red staining. Briefly, MSC-seeded well plates were fixed in 4% paraformaldehyde before incubation in 40mM Alizarin Red to detect mineralization. Histological images were captured, and quantification of staining was performed via colorimetric analysis. Briefly, wells were treated with 10% acetic acid and incubated for 30 minutes. The fluid was transferred to microcentrifuge tubes and heated to 85°C for 10 minutes. Samples were centrifuged at 20,000g for 15 minutes, supernatant was collected and resuspended in 10% ammonium hydroxide prior to reading the absorbance 405nm. Results are expressed as relative absorbance units (RAUs).

### Non-Inflammatory and Inflammatory Chondrogenic Culture Conditions

To induce chondrogenic differentiation, MSCs were seeded in pellets (1.0 x10^5^ cells per pellet; n=3 pellets/condition/cell-type) and cultured in differentiation media (StemPro Chondrogenic Media; ThermoFisher Scientific) for 28 days. For inflammatory conditions, differentiation media was supplemented with IL-1β and TNF-α at the concentrations described above. Differentiation was assessed histologically via Alcian Blue staining and quantitatively via pellet area analysis, the percent area of pellet stained positively for GAG, DNA, and GAG content analyses, respectively.

### Analysis of Chondrogenic Pellet Area

MSC pellets were imaged using phase contrast in well plates. Cross-sectional area [mm^2^] of pellets (n=3/condition/cell-type) were calculated from pellet diameters obtained using NIH ImageJ software.

### Analysis of Chondrogenic Pellet Glycosaminoglycan Staining

Chondrogenic MSC pellets (n=3/condition/cell-type) were fixed in 10% non-buffered formalin, paraffin embedded and sectioned to 5μm. Slides were stained with Alcian Blue (1% Alcian Blue in 3% aqueous acetic acid; pH 2.5) and counterstained with 0.1% aqueous Nuclear Fast Red for visualization of glycosaminoglycan deposition and cell nuclei, respectively. Histological images were captured, and the percentage of the total cell pellet area stained positively for GAG was quantified via color threshold analysis using NIH ImageJ software.

### Analysis of Chondrogenic Pellet DNA and Glycosaminoglycan Content

Chondrogenic MSC pellets (n=3/condition/cell-type) were analyzed for DNA and GAG content using PicoGreen and DMMB assays, respectively according to manufacturer’s instructions. Briefly, cell pellets were digested in PBE buffer (pH 7.5) containing 5 mM L-Cysteine, 100 mM dibasic phosphate buffer, and 5 mM EDTA and 125 μg/mL papain at 65°C for 24 hours prior to analysis via respective assays. GAG content was determined from a standard curve containing known concentrations of chondroitin-6-sulfate. DNA content was determined from a standard curve developed from known concentrations of DNA supplied by the manufacturer.

### Microscopic Imaging

All images were captured on a Zeiss Axio Vert.A1 microscope with AxioVision software (SE64 Rel. 4.9.1 SP08-2013).

### Statistics

Statistical analysis was performed using GraphPad Prism 7 software. Quantitative results are expressed as a mean ± standard error (SEM). Comparisons were performed via Student’s t-tests of equal variance comparing between control and inflammatory study groups at respective time-points. Significance was defined as p≤0.050, and significant trends were defined as p≤0.080.

## RESULTS

### Impact of Inflammation on Proliferation of AD- and AM-MSCs

In general, inflammation promoted an increase in cell number between 12- and 48-hours in both AD- and AM-MSCs compared to control conditions; however, this was found to be significant only in AD-MSC cultures. The number of AD-MSCs significantly increased over time in inflammatory media (12-hours: 113,000 ± 4,163 cells, 48-hours: 221,000 ± 8,021 cells; p<0.001), but not in control media (12-hour: 119,350 ± 22,650 cells, 48-hour: 173,333 ± 20,305 cells). The number of AM-MSCs also increased over time in inflammatory (12-hour: 87,250 ± 8,151 cells, 48-hour: 109,667 ± 5,696 cells) and control media (12-hour: 101,250 ± 2,250 cells, 48-hour: 114,500 ± 12,500 cells), however these increases were not significant.

Comparing the two MSC types, inflammation resulted in a significantly higher number of AD-MSCs at both 12- (p=0.048) and 48- (p<0.001) hours compared to AM-MSCs.

### Impact of Inflammation on Cytokine Production by AD- and AM-MSCs

Inflammation resulted in an increase in MSC production of both pro- and anti-inflammatory cytokines (Figures 1&2). Compared to controls at 12- and 48-hours, respectively, AD-MSCs in inflammation demonstrated significant increases in the production of PGE_2_, MCP-1, IL-1β and IL- 8 (Table 1). Compared to controls at 12- and 48-hours, respectively, AM-MSCs in inflammation demonstrated significant increases in the production of PGE_2_, MCP-1, IL-1β, IL-6, IL-8 and IL-10 (12-hour: 5.37 ± 0.73 pg/ml; p=0.002, 48-hour: 14.47 ± 2.39 pg/ml; p=0.018) (Table 1).

**Fig1.**
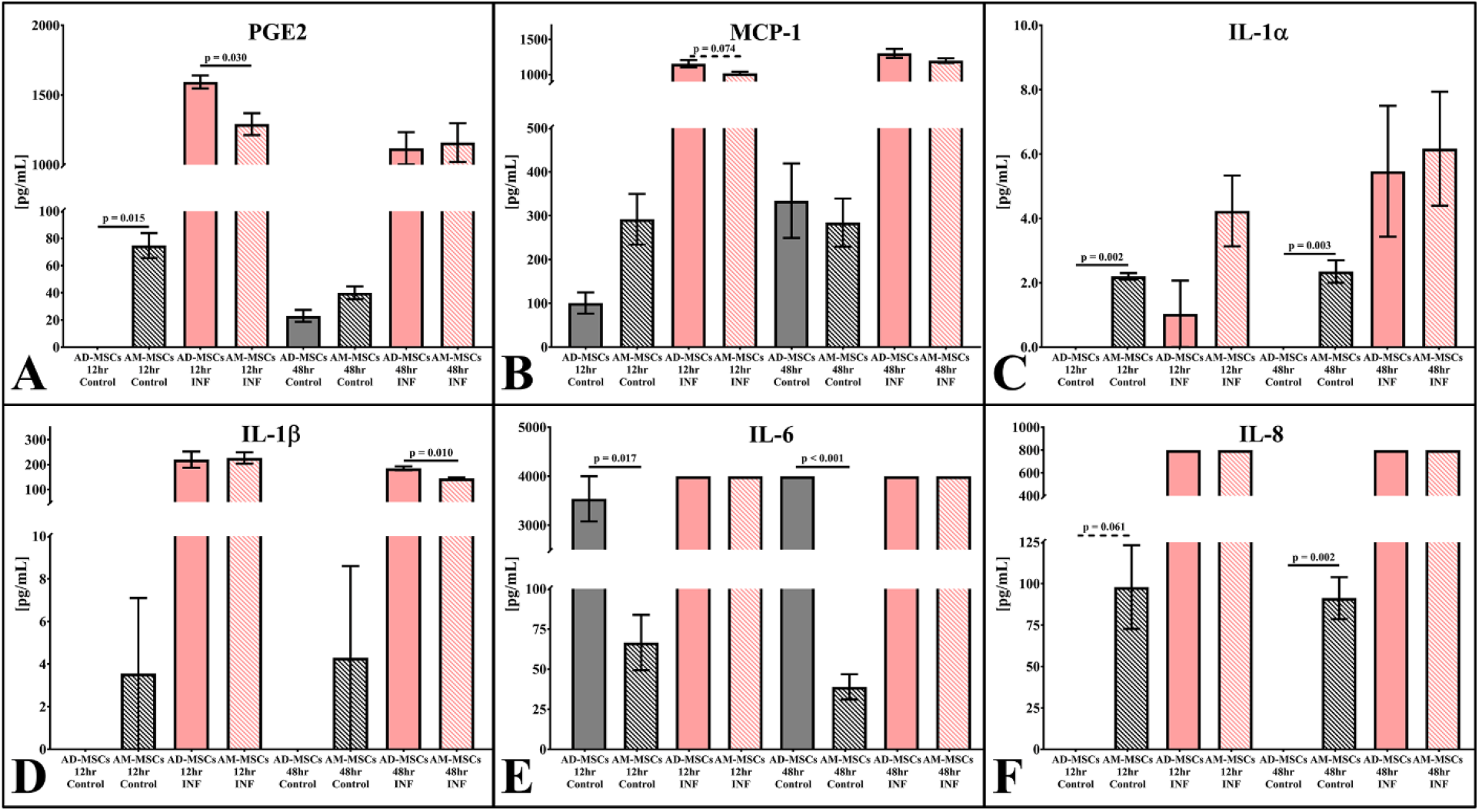
Pro-inflammatory cytokine production profile of human MSCs in control and inflammatory culture conditions. Graphs depicting production of pro-inflammatory cytokines by AD- and AM-MSCs following 24- and 48-hours of culture in the absence (Control; gray bars) or presence of IL-1β and TNF-α (INF: pink bars). Solid lines connecting bars indicate significant differences between cell types within the same culture condition (p≤0.050). Dotted lines connecting bars indicate a significant trend between cell types within the same culture condition (p≤0.080).

**Fig2.**
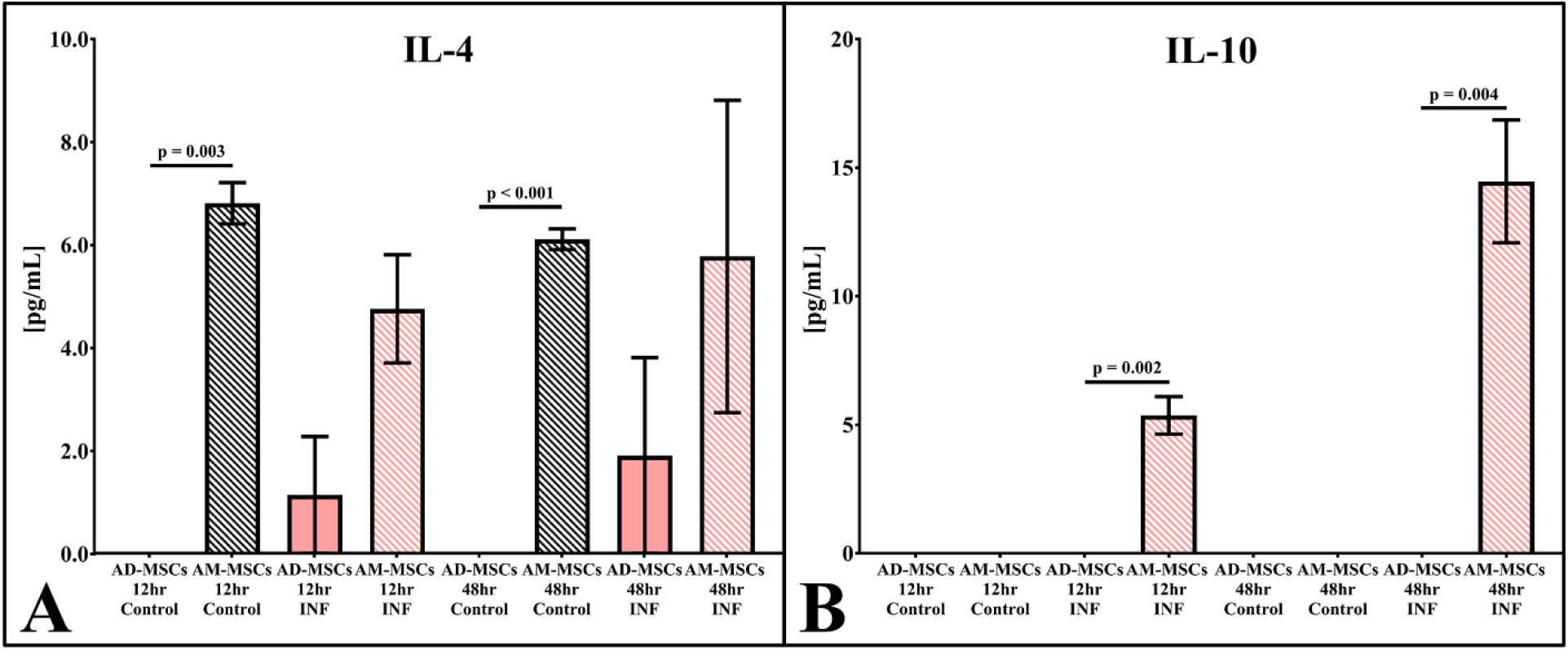
Anti-inflammatory cytokine production profile of human MSCs in control and inflammatory culture conditions. Graphs depicting production of anti-inflammatory cytokines by AD- and AM-MSCs following 24- and 48-hours of culture in the absence (Control; gray bars) or presence of IL-1β and TNF-α (INF: pink bars). Solid lines connecting bars indicate significant differences between cell types within the same culture condition (p≤0.050). Dotted lines connecting bars indicate a significant trend between cell types within the same culture condition (p≤0.080).

**Table 1.** Summary of cytokine production profile of human MSCs in control and inflammatory culture conditions. Table depicting pro- and anti-inflammatory cytokine concentrations produced by AD- and AM-MSCs following 24- and 48-hours of culture in the absence (Control) or presence (INF) of IL-1β and TNF-α. * indicates a significant difference between culture conditions within MSC type (p≤0.050).

| [pg/mL] | AD-MSCs |  |  |  |  |  |
| --- | --- | --- | --- | --- | --- | --- |
|  | 12hr |  |  | 48hr |  |  |
|  | Control | INF | p-value | Control | INF | p-value |
| **PGE2** | 0.00 ± 0.00 | **1593.80 ± 46.51\*** | **<0.001** | 23.00 ± 4.38 | **1118.30 ± 115.56\*** | **0.005** |
| **MCP-1** | 100.60 ± 24.30 | **1154.20 ± 52.57\*** | **<0.001** | 334.27 ± 85.05 | **1301.70 ± 63.71\*** | **<0.001** |
| **IL-1α** | 0.00 ± 0.00 | 1.03 ± 1.03 | 0.495 | 0.00 ± 0.00 | 5.47 ± 2.03 | 0.054 |
| **IL-1β** | 0.00 ± 0.00 | **220.27 ± 32.48\*** | **0.013** | 0.00 ± 0.00 | **185.40 ± 7.63\*** | **<0.001** |
| **IL-4** | 0.00 ± 0.00 | 1.13 ± 1.13 | 0.495 | 0.00 ± 0.00 | 1.90 ± 1.90 | 0.370 |
| **IL-6** | 3537.80 ± 462.20 | 4000.00 ± 0.00 | 0.272 | 4000.00 ± 0.00 | 4000.00 ± 0.00 | 0.999 |
| **IL-8** | 0.00 ± 0.00 | **800.00 ± 0.00\*** | **<0.001** | 0.00 ± 0.00 | **800.00 ± 0.00\*** | **<0.001** |
| **IL-10** | 0.00 ± 0.00 | 0.00 ± 0.00 | 0.999 | 0.00 ± 0.00 | 0.00 ± 0.00 | 0.999 |

| [pg/mL] | AM-MSCs |  |  |  |  |  |
| --- | --- | --- | --- | --- | --- | --- |
|  | 12hr |  |  | 48hr |  |  |
|  | Control | INF | p-value | Control | INF | p-value |
| **PGE2** | 74.72 ± 9.17 | **1291.40 ± 78.47\*** | **0.001** | 39.99 ± 4.71 | **1158.70 ± 138.99\*** | **0.008** |
| **MCP-1** | 291.85 ± 57.85 | **1018.30 ± 22.34\*** | **<0.001** | 284.25 ± 54.95 | **1198.60 ± 32.20\*** | **<0.001** |
| **IL-1α** | 2.20 ± 0.10 | 4.23 ± 1.09 | 0.248 | 2.35 ± 0.35 | 6.17 ± 1.77 | 0.196 |
| **IL-1β** | 3.55 ± 3.55 | **226.73 ± 22.81\*** | **0.005** | 4.30 ± 4.30 | **144.10 ± 4.57\*** | **<0.001** |
| **IL-4** | 6.80 ± 0.40 | 4.75 ± 1.05 | 0.210 | 6.10 ± 0.20 | 5.77 ± 3.03 | 0.938 |
| **IL-6** | 66.55 ± 17.25 | **4000.00 ± 0.00\*** | **<0.001** | 38.95 ± 7.85 | **4000.00 ± 0.00\*** | **<0.001** |
| **IL-8** | 97.95 ± 25.25 | **800.00 ± 0.00\*** | **<0.001** | 91.25 ± 12.65 | **800.00 ± 0.00\*** | **<0.001** |
| **IL-10** | 0.00 ± 0.00 | **5.37 ± 0.73\*** | **0.002** | 0.00 ± 0.00 | **14.47 ± 2.39\*** | 0.018 |

*\* indicates significance (p≤0.050) compared between culture conditions at respective time point*

Comparing the two MSC types, AD-MSCs produced significantly more PGE_2_ and IL-1β at 12- and 48-hours, respectively compared to AM-MSCs in inflammatory media (Figure 1). Additionally, AD-MSCs produced more MCP-1 (p=0.074) at 12-hours compared to AM-MSCs in inflammatory media (Figure 1). Conversely, AM-MSCs produced significantly more IL-10 at 12- (p=0.002) and 48-hours (p=0.004) compared to AD-MSCs when cultured in inflammatory media (Figure 2). Of note, significant differences were also observed comparing the cytokine production of AD-MSCs and AM-MSCs in control media (Figures 1&2, Table 1).

### Effect of Inflammation on Extracellular Matrix Mineralization by AD- and AM-MSCs

In non-inflammatory conditions, histological imaging confirmed that AD- and AM-MSCs produced mineralized ECM as evidenced by positive (red) alizarin staining on the bottom of all wells (Figure 3). Additionally, macroscopic images of wells illustrated the formation of white, mineralized ECM nodules in AD- and AM-MSC cultures (Figure 3; inserts). Inflammation affected osteogenesis of both MSC types to different degrees. Inflammation resulted in a significant reduction in the number of mineralized nodules (1.33 ± 0.04 per well; p=0.013) in AD- MSC cultures compared to control conditions (3.33 ± 0.33 per well). However, inflammation did not significantly affect nodule diameter (inflammation: 0.22 ± 0.33mm, control: 0.18 ± 0.02mm) or the amount of Alizarin Red staining (inflammation: 3.34 ± 0.05 RAU, control 3.46 ± 0.07 RAU) in AD-MSC cultures (Figure 3). Conversely, inflammation resulted in a significant increase the number of nodules (inflammation: 27.67 ± 0.23 per well, control: 9.33 ± 1.86 per well; p=0.009) in AM-MSC cultures; however, a significant reduction in nodule diameter (inflammation: 0.23 ± 0.02mm, control: 0.36 ± 0.05mm; p=0.002) and the amount of Alizarin Red staining (inflammation: 1.08 ± 0.06 RAU, control: 1.80 ± 0.03 RAU; p<0.001) was observed in AM-MSC cultures (Figure 3).

**Fig3.**
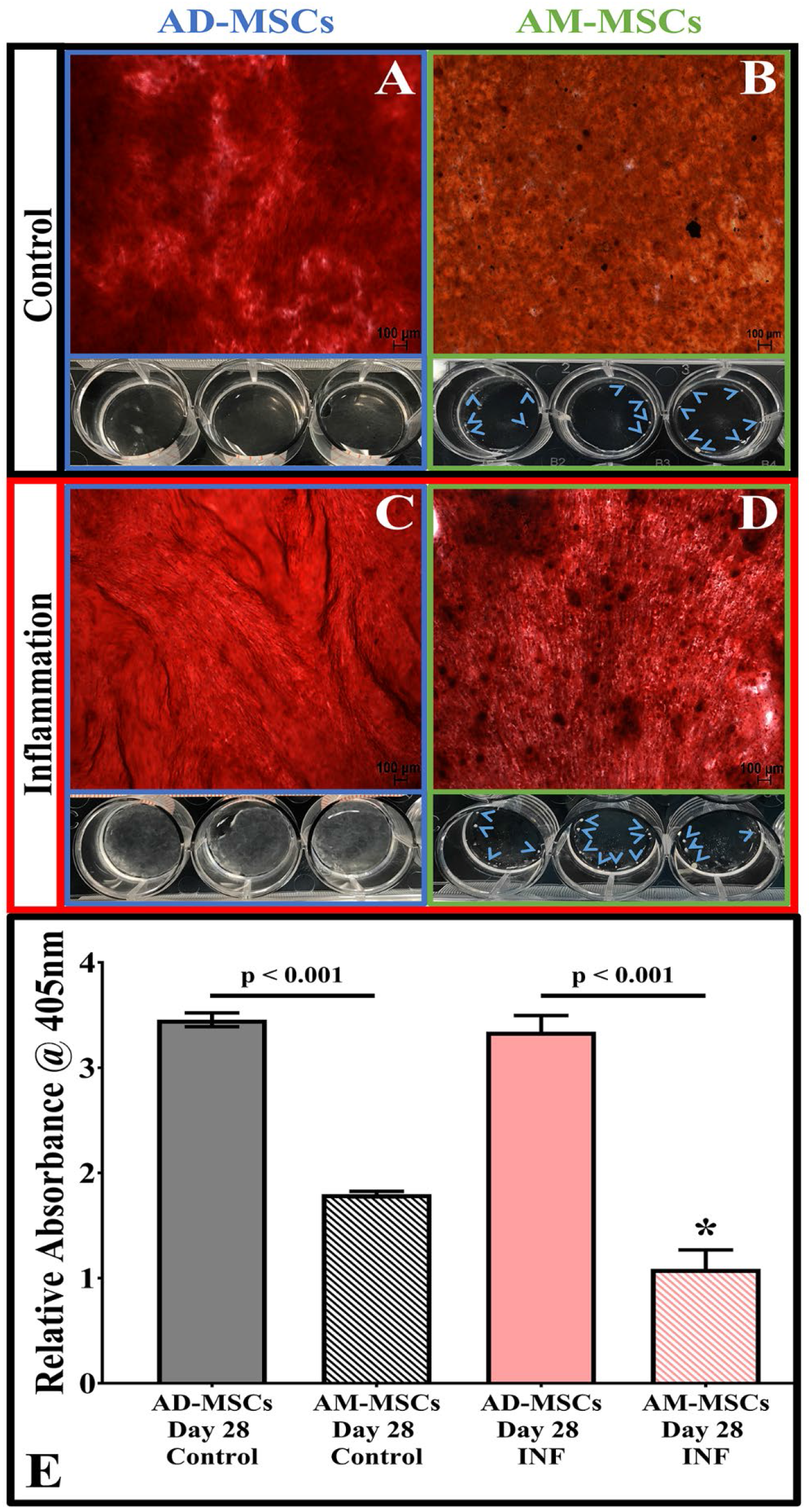
Osteogenic potential of human MSCs in control and inflammatory culture conditions. Representative alizarin red staining (red = mineralized ECM) in **A)** AD- and **B)** AM-MSC osteogenic cultures in control (non-inflammatory) conditions. (inserts = macroscopic imaging of wells illustrating mineralization (white pellets indicated by arrowheads). Representative alizarin red staining (red = mineralized ECM) in **C)** AD- and **D)** AM-MSC osteogenic cultures in inflammatory conditions. (inserts = macroscopic imaging of wells illustrating mineralization (white pellets). **E)** Graph depicting quantification of alizarin red staining of AD- and AM-MSCs in control (non-inflammatory) and inflammatory conditions. Magnification = 50x. Scale bars = 100μm. Solid lines connecting bars indicate significant differences between MSC type within the same culture conditions (p≤0.050). * Indicates significant difference between culture conditions within MSC type (p≤0.050).

Comparing the two MSC types, AD-MSC osteogenesis was impacted to a lesser degree by the presence of inflammation as compared to AM-MSCs. AD-MSCs demonstrated the formation of a thin white film of ECM coating the bottom of the wells resulting in significantly fewer nodules (p=0.001) compared to AM-MSC cultures in inflammatory differentiation media (Figure 3; inserts). However, AD-MSC cultures had significantly higher amounts of total alizarin red staining (p<0.001) compared to AM-MSC cultures in inflammatory differentiation media (Figure 3).

### Effect of Inflammation on Glycosaminoglycan-Containing Extracellular Matrix Formation by AD- and AM-MSCs

In non-inflammatory conditions, histological imaging confirmed that AD- and AM-MSCs produced ECM containing GAG as evidenced by positive (blue) staining within all pellets (Figure 4). Inflammation did affect chondrogenesis of both MSC types to different degrees. Inflammation resulted in ECM irregularities and voids within the AD-MSC pellets which appeared more fibrous (Figure 4) and demonstrated significant reductions in the percentage of pellet area staining positively for Alcian blue (inflammation: 34.75 ± 2.49%, control: 57.83 ± 2.10%; p<0.001) and overall pellet cross-sectional area (inflammation: 2.76 ± 0.18mm^2^, control: 4.25 ± 0.22mm^2^; p=0.006) compared to control conditions (Figure 4). Conversely, AM-MSC pellets remained intact with no voids under inflammatory conditions (Figure 4) and did not demonstrate a significant reduction in pellet cross-sectional area (inflammation: 5.67 ± 0.26mm^2^, control: 6.20 ± 0.61mm^2^) (Figure 4). However, the percentage of pellet area staining positively for Alcian blue (inflammation: 82.03 ± 3.26%, control: 92.86 ± 1.01; p=0.019) was significantly reduced.

**Fig4.**
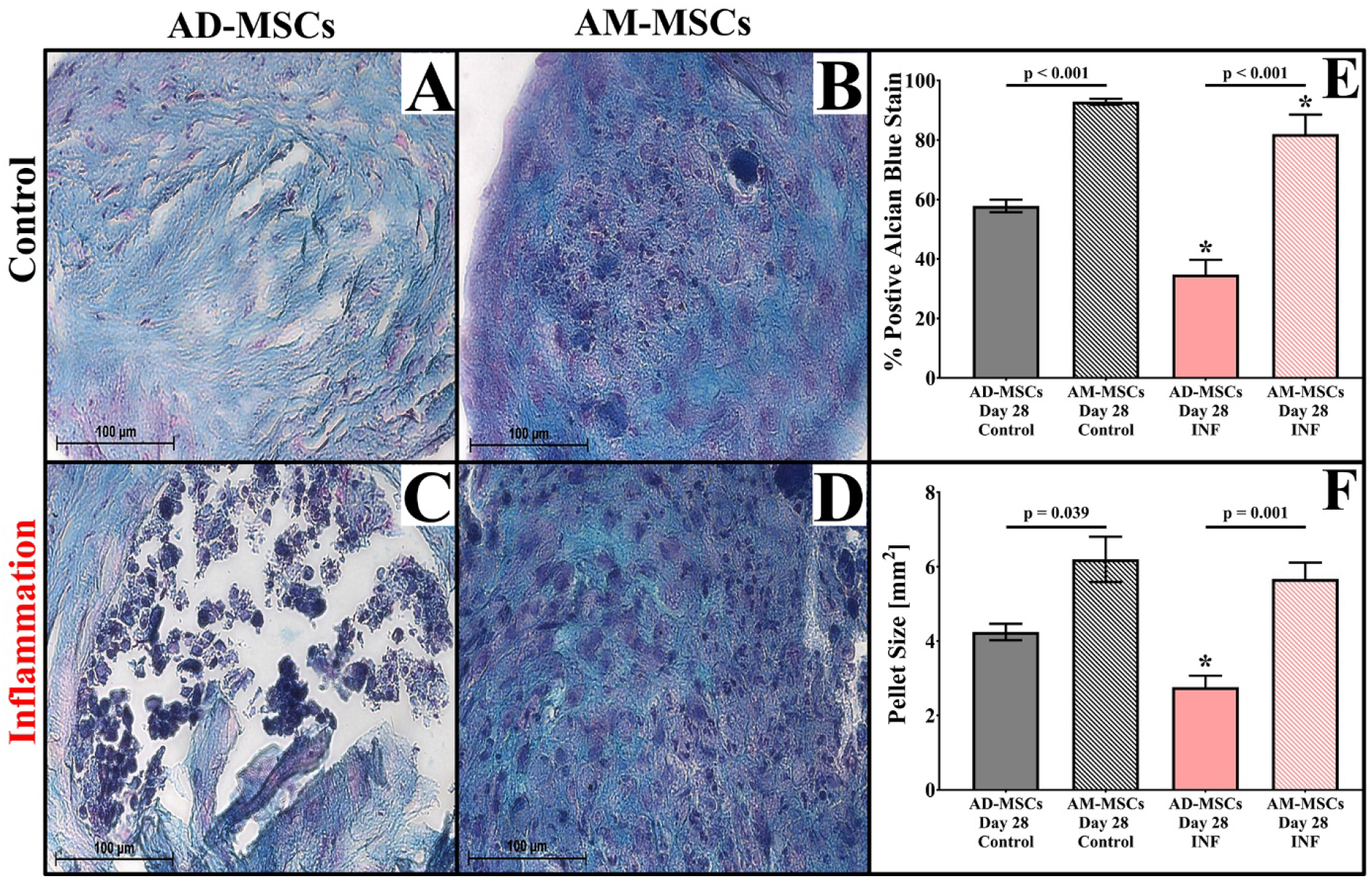
Chondrogenic potential of human MSCs in control and inflammatory culture conditions. Representative Alcian blue staining (blue = GAG-containing ECM, purple = cell nuclei) in **A)** AD- and **B)** AM-MSC chondrogenic pellet cultures in control (non-inflammatory) and **C)** AD- and **D)** AM-MSC chondrogenic pellet cultures in inflammatory conditions. Graphs depicting quantification of **E)** the percentage of pellet area stained positively for GAG and **F)** cross-sectional pellet area of AD- and AM-MSCs in control (non-inflammatory) and inflammatory conditions. Magnification = 400x. Scale bars = 100μm. Solid lines connecting bars indicate significant differences between MSC type within the same culture conditions (p≤0.050). * Indicates significant difference between culture conditions within MSC type.

Comparing the two MSC types, AM-MSC chondrogenesis was impacted to a lesser degree by the presence of inflammation as compared to AD-MSCs. AM-MSCs had significantly greater GAG staining (p<0.001) and cross-sectional pellet area (p=0.04) compared to AD-MSCs in inflammation.

### Effect of Inflammation on DNA and Glycosaminoglycan Quantification by AD- and AM-MSCs

Inflammation resulted in increased DNA and GAG content in both AD- and AM-MSC pellets (Figure 5). AD-MSC pellet DNA (inflammation: 0.14 ± 0.01 μg/pellet, control: 0.05 ± 0.01 μg/pellet; p=0.002) and GAG content (inflammation: 5.90 ± 0.57 μg/pellet, control: 2.63 ± 0.40 μg/pellet; p=0.009) significantly increased compared to control conditions (Figure 5). AM-MSC pellet DNA (inflammation: 0.18 ± 0.03 μg/pellet, control: 0.10 ± 0.001 μg/pellet; p=0.048) and GAG content (inflammation: 8.34 ± 0.88 μg/pellet, control: 6.95 ± 0.25 μg/pellet) also increased, but the latter was not significant (Figure 5). However, normalization of pellet GAG content to DNA indicated that inflammation significantly hindered GAG production by AD-MSCs (41.46 ± 1.46 μg GAG/ μg DNA) and AM-MSCs (47.99 ± 2.66 μg GAG/ μg DNA) compared to controls (AD-MSCs: 49.64 ± 1.36 μg GAG/ μg DNA, AM-MSCs: 68.78 ± 3.45 μg GAG/ μg DNA) (Figure 5). Comparing the two MSC types, AM-MSCs produced higher GAG content compared to AD-MSCs in inflammation (p=0.08).

**Fig5.**
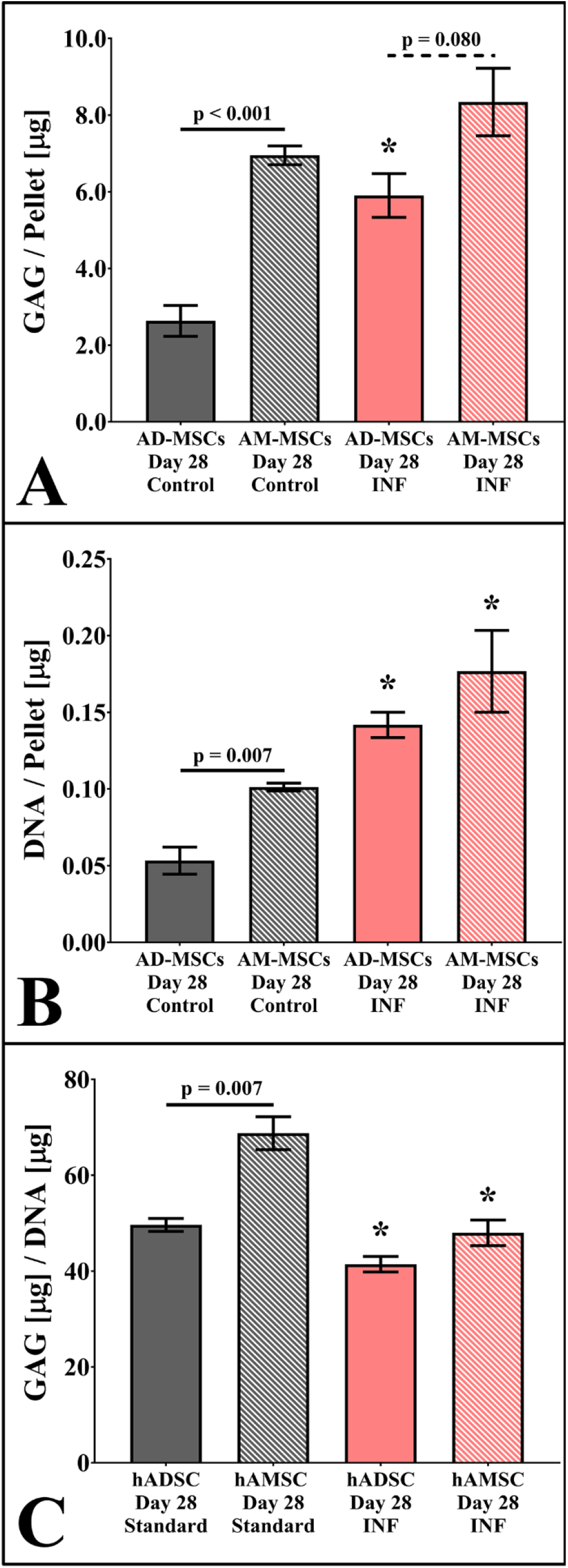
DNA and GAG quantification of human MSCs in control and inflammatory culture conditions. Graphs depicting quantification of **A)** GAG and **B)** DNA content per AD- and AM- MSC chondrogenic pellet culture in control (non-inflammatory) and inflammatory conditions. **C)** Graph depicting GAG normalized to DNA content for AD- and AM-MSC chondrogenic pellet culture in control (non-inflammatory) and inflammatory conditions. Solid lines connecting bars indicate significant differences between MSC type within the same culture conditions (p≤0.050). * Indicates significant difference between culture conditions within MSC type. Dotted lines connecting bars indicate a significant trend between cell types within the same culture condition (p≤0.080).

## DISCUSSION

Our findings suggest that inflammation affects cytokine production and the differentiation capacity of AD- and AM-MSCs differently. More specifically, AM-MSCs produced more anti-inflammatory cytokines and cartilaginous ECM compared to AD-MSCs when cultured in the presence of inflammation. Conversely, AD-MSC culture in inflammation resulted in increased proliferation and production of more pro-inflammatory cytokines and mineralized ECM compared to AM-MSCs. These differences may have significant clinical implications when considered in the context of the local environment of the degenerate IVD.

The first significant difference comparing the two MSC sources was that culture in inflammation resulted in a significant increase in AD-MSC numbers compared to AM-MSCs. In many tissues, inflammation has been shown to promote activation, migration, and proliferation of resident MSCs as a mechanism to increase cell numbers at a site of injury and prime them to promote tissue repair and regeneration. [6,17] Thus, increases in MSC number is generally thought to be a beneficial response in normal wound healing conditions. However, in the context of the degenerate IVD, it has been hypothesized that an increase in MSC numbers could quickly deplete the already limited nutrient supply within the tissue and thus may not provide therapeutic benefit.[18] Furthermore, it is essential to identify the species and amounts of soluble cytokines being released by these MSCs and to consider their effects on other local cells present in the degenerate IVD.

Thus, the second significant difference found comparing the two MSC sources was that AD-MSCs produced more PGE_2_ and MCP-1 at the early (12-hour) time-point compared to AM-MSCs. MCP-1 is produced by many cell types to recruit macrophages, and other inflammatory cells to sites of tissue injury, inflammation, or infection.[18] PGE_2_ produced by MSCs is thought to exert an anti-inflammatory effect by shifting infiltrating pro-inflammatory, M1-macrophages towards a pro-regenerative, M2 phenotype.[5] Thus, enhanced recruitment of macrophages via increased expression of MCP-1 coupled with increased expression of PGE_2_ could subsequently quell local tissue inflammation. However, in the context of the intact degenerate IVD relatively few blood vessels are found, and thus infiltration of exogenous macrophages is limited to only the periphery of the IVD (e.g. cartilaginous end-plates). [19] Therefore, increased concentrations of MCP-1 and PGE_2_, as observed by AD-MSCs in the current study, may not be beneficial in the context of the degenerate IVD. More specifically, studies have demonstrated that PGE_2_ has a negative impact on IVD cell health and ECM homeostasis even after short-term exposure.[20,21] Moreover, increased MCP-1 expression by cells in the IVD correlate positively with increasing histological grades of IVD degeneration.[3] However, additional studies investigating the impact of elevated levels of MCP-1 on IVD cell-mediated ECM homeostasis in a three-dimensional (e.g., IVD tissue mimetic) inflammatory environment are warranted.[22]

Another significant difference identified when comparing the cytokine production of AD- and AM-MSCs in the presence of inflammation was that AM-MSCs produce significantly less pro-inflammatory IL-1β and significantly more anti-inflammatory IL-10 at the later time-point (48-hour) compared to AD-MSCs. IL-1β is a pro-inflammatory cytokine, which along with TNF-α, plays a key role in the progression of IVD degeneration.[3] Increased concentrations of IL-1β have been shown to cause IVD cells to increase production of tissue degrading proteases, pro-inflammatory and chemotactic cytokines, concomitant with inhibiting ECM biosynthesis.[2,3] Thus, the increased production of IL-1β by AD-MSCs observed herein suggests that this MSC type could exacerbate the pro-inflammatory environment more than AM-MSCs. Conversely, IL-10 is a potent anti-inflammatory and immune suppressant which plays a prominent autoregulatory role in the production of pro-inflammatory cytokines by macrophages. [23] In the context of the degenerate IVD, IL-10 is found in increased levels compared to non-degenerate tissues, [16] suggesting an endogenous attempt to quell inflammation. Moreover, administration of exogenous IL-10 to degenerate IVD cell cultures has been shown to decrease transcription of TNF-α and IL-1β. [24] Thus, the observed increase in production of IL-10 by AM-MSCs in the inflammatory conditions studied herein could serve as an exogenous source of anti-inflammatories which were not found to be produced by AD-MSCs. Of note, the decreased production of IL-1β observed in the AM-MSC group may be related to their increased production of IL-10 suggesting the possibility of a similar autoregulatory mechanism to that observed in macrophages, however further study of the specific mechanisms involved are warranted.

Another key difference observed comparing MSC types was that the presence of inflammation detrimentally impacted chondrogenesis of AD-MSCs to a greater extent compared to AM-MSCs. It is well established that the presence of TNF-α and/or IL-1β impairs chondrogenesis of MSCs. This is mediated in part via increased translocation of nuclear factor kappa beta (NF-kβ) and inhibition of the transcriptional activator sex determining region Y-box 9 (Sox-9) and transforming growth factor beta (TGF-β) signaling which is required for chondrogenesis. [25,26] In the present study, although chondrogenesis appeared to be hampered by inflammation in both MSC types compared to their respective non-inflammatory controls, AM-MSC cultures yielded larger, intact chondrogenic pellets, enhanced histological staining, and quantification of GAG compared to AD-MSCs. Similar differences were observed by others when comparing the effects of IL-1β (10ng/ml), TNF-α (50ng/ml), or human osteoarthritic synovial fluid on AD- and BM-MSC chondrogenesis.[27] In the previous studies, it was found that although AD-MSCs pellets demonstrated histological irregularities including; voids in the pellet, altered cell nucleus morphology, smaller pellet size, and reduced GAG staining compared to non-inflammatory controls, they appeared to fare better than BM-MSCs cultures.[27] The reasons for the observed differences between these studies are still unclear and require further investigation. However, it could be hypothesized that differences in MSC cytokine receptor expression and thus sensitivity to inflammation may play a role. Additionally, it is possible that AD-MSCs could have proliferated more in inflammatory pellet cultures compared to AM-MSCs resulting in a higher nutrient demand and thus resulting in the formation of a necrotic core and reduction in pellet GAG staining. Although there was no significant difference found in pellet DNA content comparing AD- and AM-MSCs at the study end-point, further analysis to confirm this hypothesis is warranted. Regarding the clinical application of MSCs for IVD degeneration, our results suggest that AM-MSCs may have an enhanced ability to produce ECM containing GAG, as is typically found within the nucleus pulposus (NP)-region of the IVD, compared to AD-MSCs. Thus, AM-MSCs may have a higher propensity to regenerate IVD tissue in mild- to moderately degenerate IVDs.

The final significant difference observed comparing MSC types was that the presence of inflammation detrimentally impacted the osteogenic differentiation of AM-MSCs, but not AD-MSCs. Previous *in vitro* studies have demonstrated that inflammatory mediators TNF-α and IL-1β enhance osteogenesis of human MSCs in a dose-dependent manner,[28] but only if the MSCs are osteogenically primed or are cultured in the presence of osteogenic signals.[29] While we observed similar effects in AD-MSC cultures (as was indicated macroscopically by more uniform mineralization in the wells compared to their non-inflammatory conditions), the converse was true for AM-MSCs. The difference could be explained in part by the short-term cytokine data which not only demonstrated elevated levels of IL-1β produced by AD-MSC cultures but also higher concentrations of PGE_2_. PGE_2_ has been shown to be a key cytokine necessary for promoting bone healing and regeneration. [30,31] However, to further confirm this hypothesis cytokine analysis on the inflammatory differentiation media at early- and late time-points are warranted. From a clinical perspective, these results suggest that AD-MSCs may be more prone to forming bone in the context of the inflammatory environment of the degenerate IVD compared to AM-MSCs. Provided an osteoinductive cue (e.g., autologous bone graft), AD-MSCs may be more amenable to promoting fusion to combat the effects of severe degeneration and pseudarthrosis as compared to AM-MSCs. Conversely, AM-MSCs may better support IVD soft-tissue regeneration.

As with any study, limitations were noted. First, the effect of other degenerate IVD environmental stimuli including other inflammatory cytokines, oxygen tension, pH, limited nutrient supply and osmolarity, which have been shown to impact MSC effector function and viability, were not investigated. [32] For example, studies have shown that MSC metabolism is highly dependent upon the presence of glucose, [33] and that MSCs may be tolerant of or adapt to nutrient deprivation. [34,35] Moreover, hypoxia may enhance MSC differentiation towards a IVD cell-like phenotype. [36, 37] However, when combined with hypoxia, long-term nutrient deprivation has been shown to lead to MSC death. [38] Considering this information, future studies comparing AD- and AM-MSCs survival and effector function should be performed in an environment reminiscent of nutrient and oxygen deprivation as well as inflammation and extracellular matrix mechanical strain as would be expected in the degenerate IVD. Completion of such studies would help to provide a more definitive answer as to which MSC source may be more amenable for therapeutic use. Regardless, our objective herein was to elucidate the effects of the primary inflammatory mediators involved in IVD degeneration without overcomplicating the study system. Secondly, we did not specifically investigate the effect of inflammation on MSC differentiation toward IVD cell-specific phenotypes (e.g., nucleus pulposus and annulus fibrosus cells). However, identifying distinguishing phenotypic markers for IVD cells continues to be investigated, [39–41] and repeatable differentiation protocols to achieve these phenotypes are still being defined. [42] Moreover, we chose to evaluate bone and cartilage formation as these phenotypes are relevant in the context of the degenerate IVD. For example, the ectopic bone formation has been observed following leakage of intradiscally injected MSCs from degenerate IVDs,[43] and products containing MSCs have been investigated to improve IVD fusions.[44,45] Additionally, although the relative quantities are different, the primary ECM components that comprise the nucleus pulposus region of the IVD are similar to that of articular cartilage.[46] Another limitation to the study was that the differential response of the MSC types to the same inflammatory conditions could have been due to differences in MSC age and donor variability which was not controlled for in this study. However, it was not feasible to obtain amnion and adipose tissue samples from the same donors, and moreover, the younger age of amnion-derived MSCs may represent an actual clinical advantage over other adult sources. [47]

## CONCLUSION

AM-MSCs may be more amenable to promote IVD tissue regeneration as compared to AD-MSCs in the context of the inflammatory environment found in the degenerate IVD. Conversely, AD-MSCs may be more prone to form bone and thus promote IVD fusion. Although our studies did not compare AD- and AM-MSC survival in a complex environment reminiscent of nutrient and oxygen deprivation as well as mechanical strains expected in the degenerate IVD, the results herein provide the impetus for further investigation into the mechanisms underlying the observed differences between AD- and AM-MSC efficacy as therapeutics for IVD degeneration.

## ACKNOWLEDGEMENTS

Research reported in this publication was supported in part by the National Institute of General Medical Sciences of the National Institutes of Health (5P20GM103444). RB is supported by the National Science Foundation Graduate Research Fellowship (Award number: 2011382). Disclosures: **RB, AM, MB, JM:** No competing financial interests exist. **SG:** Speaking and/or Teaching Arrangements-OrthoFix. **JT:** Royalties-Arthrex; Consulting-Arthrex; Speaking and/or Teaching Arrangements-Arthrex and Johnson & Johnson; Grants-Mayo Clinic.

---

**Supplemental Table 1.** Summary of osteogenesis and chondrogenesis differentiation quantification of human MSCs in control and inflammatory culture conditions. Table depicts the representative quantitative outcomes of the differentiation potential of AD- and AM-MSCs following 28 days of culture in the absence (Control) or presence (INF) of IL-1β and TNF-α. Osteogenesis outcomes include: Relative absorbance units (RAU) of Alizarin Red staining for mineralized extracellular matrix production. Chondrogenesis outcomes include: Normalized glycosaminoglycan (GAG) content to the respective pellet’s DNA content, percentage of pellet positively stained for Alcian Blue indicating GAG presence, pellet cross-sectional area, and percent reduction of pellet cross-sectional area from its respective Day 0 measurements. * indicates a significant difference between culture conditions within MSC type (p≤0.050).

| Day 28 | AD-MSCs |  | AM-MSCs |  |
| --- | --- | --- | --- | --- |
|  | Control | INF | Control | INF |
| Alizarin Red staining for Osteogenesis [RAU] | 3.46 ± 0.07 | 3.34 ± 0.05 | 1.80 ± 0.03 | **1.09 ± 0.06\*** |
| Normalized GAG to DNA [μg GAG / μg DNA] | 49.64 ± 1.36 | **41.46 ± 1.62\*** | 68.78 ± 3.45 | **47.99 ± 2.66\*** |
| Alcian Blue Staining for GAG [% of Pellet Positively Stained] | 57.83 ± 2.10 | **34.75 ± 2.49\*** | 92.86 ± 1.01 | **82.03 ± 3.26\*** |
| Pellet Cross-Sectional Area [$\text{mm}^2$] | 4.25 ± 0.22 | **2.76 ± 0.18\*** | 6.20 ± 0.61 | 5.67 ± 0.26 |
| % Reduction of Pellet Cross-Sectional Area from Day 0 | 5.63 ± 10.62 | **38.59 ± 7.66\*** | 8.14 ± 1.86 | 19.42 ± 7.03 |

*\* indicates significance ( $p \leq 0.050$ ) compared between culture conditions*

**Author contribution statement**
RB, SG, JT, and JM contributed to conception and design of the study. RB, AM, MB, JM contributed to data acquisition, analysis, and data interpretation. RB and JM contributed to drafting of the manuscript and statistical analysis. SG, JT, and JM provided administrative, technical, and material support for this study. All authors provided critical revision of the manuscript for important intellectual content, and all have read and approved the final submitted manuscript.

